# Low-pH induced reversible reorganizations of chloroplast thylakoid membranes - as revealed by small-angle neutron scattering

**DOI:** 10.1101/069211

**Authors:** Renáta Ünnep, Ottó Zsiros, Zsolt Hörcsik, Márton Markó, Anjana Jajoo, Joachim Kohlbrecher, Győző Garab, Gergely Nagy

## Abstract

Energization of thylakoid membranes brings about the acidification of the lumenal aqueous phase, which activates important regulatory mechanisms. Earlier Jajoo and coworkers (2014 FEBS Lett. 588:970) have shown that low pH in isolated plant thylakoid membranes induces changes in the excitation energy distribution between the two photosystems. In order to elucidate the structural background of these changes, we used small-angle neutron scattering on thylakoid membranes exposed to low p^2^H and show that gradually lowering the p^2^H from 8.0 to 5.0 causes small but well discernible reversible diminishment of the periodic order and the lamellar repeat distance and an increased mosaicity – similar to the effects elicited by light-induced acidification of the lumen. Our data strongly suggest that thylakoids dynamically respond to the membrane energization and actively participate in different regulatory mechanisms.

**Highlights:**

1. Thylakoid membranes exposed to low p^2^H studied by small-angle neutron scattering
2. Acidification causes reversible shrinkage and diminished lamellar order
3. SANS changes induced by low pH resemble those due to light-induced lumenal acidification

**Abbreviations:**

NPQ: non-photochemical quenching
qE: the energy-dependent component of NPQ
Δμ_H_^+^: transmembrane electrochemical potential gradient
PSI: photosystem I
PSII: photosystem II
LET: linear electron transport
CD: circular dichroism
SANS: small-angle neutron scattering
q: scattering vector or momentumtransfer
I: intensity
q^*^: center position of the Bragg peak
RD: repeat distance
φ: azimuthal angle
I(φ): angular dependency of the scattering intensity
FWHM: full width at half maximum

## Introduction

In photosynthesis, charge separation followed by vectorial electron transport, coupled to proton translocation processes, creates a transmembrane electrochemical potential gradient, Δμ_H_^+^, between the inner and outer aqueous phases of the photosynthetic membranes - in chloroplasts, the lumenal and stromal sides of the thylakoid membranes. This proton-motive force, which is utilized for the synthesis of ATP, consists of an electrical field and a ΔpH component of ∼10^5^ V cm^-1^ and ∼2-3 pH units, respectively, and modulates the electron transport rate via various feedback regulatory mechanisms. The transmembrane electric potential gradient is required for metabolite and protein transport across the membranes [1]. The ΔpH component is involved, perhaps most prominently, in the photoprotective mechanisms of non-photochemical quenching (NPQ) of the first singlet excited state of chlorophyll-a [2]. In particular, qE, the energy-dependent component of NPQ depends on the acidification of the lumen [3]. It is generally agreed that NPQ requires the structural flexibility of thylakoid membranes; in fact, there are several reports demonstrating the involvement of structural changes at different levels of structural complexity [4-14]. Some of these changes might directly be linked to the generation of ΔpH, e.g. via the redistribution of ions in the ‘electrolyte’ following the generation of Δμ_H_^+^ [15, 16] and, in particular, upon the acidification of lumen and the binding of protons to different polypeptide residues [2, 17, 18].

In general, the effects of pH on many physiological processes in plants are well established and significant work has been done to explore the effects of pH on different photosynthetic processes. In addition to the involvement of lumenal acidification in NPQ mechanisms, acidic lumen leads to inhibition of Photosystem II (PSII) activity because of a reversible dissociation of Ca^2+^ from the water splitting enzyme [19]. In vitro, the oxygen evolving complex loses Ca^2+^ at pH<6.0, inhibiting water splitting and rendering the PSII reaction center inactive [20]. The linear electron transport (LET) can also be down-regulated via the back-pressure due to the build up of ΔpH [21]. The photosynthetic machinery in plants is endowed with a strong ΔpH-dependent control mechanism of LET from cytochrome b6f to PSI. By using pgr5 mutant of Arabidopsis which is deficient in strong light-induced ΔpH, it has been shown that PSI takes a central role in excess energy dissipation and control of LET [22].

In earlier works, structural and functional changes have been shown to be induced by exposing isolated thylakoid membranes to low pH [23-25]. It has been shown mainly by 77K fluorescence excitation and emission spectroscopy of isolated spinach thylakoid membranes that low pH induces a redistribution of the excitation energy between the two photosystems. By analysing data obtained on state transition and NPQ mutants of Arabidopsis it has been shown that the increase in the 77K emission of PSI and the concomitant quenching of PSII fluorescence in thylakoid membranes which were exposed to low pH cannot be accounted for by state transitions but originates from a PsbS-protonation dependent spillover of the excitation energy from PSII to PSI. It has also been shown, by using circular dichroism (CD) spectroscopy, that exposing isolated pea thylakoid membranes to low pH induces substantial but essentially fully reversible changes in the chiral macroorganization of the protein complexes, without noticeable changes in the excitonic interactions, i.e. at the level of bulk pigment-protein complexes [23]. These reorganizations in the CD were similar to those induced by light [26-28]. Here, in order to obtain more information on the nature of these membrane reorganizations, by using small-angle neutron scattering (SANS), we investigated the effect of low pH on the multilamellar organization of the thylakoid membrane system of isolated pea thylakoid membranes. Our data reveal small (≤ 2 nm) but well discernible low-pH induced shrinkage in the repeat distance of the grana thylakoid membranes and a diminishment in their periodic order, accompanied by an increased mosaicity of membranes.

## Materials and Methods

### Isolation of thylakoid membranes

Thylakoid membranes were isolated as described earlier [29] from freshly harvested three-weeks-old pea leaves (*Pisum sativum*, Rajnai törpe) grown in a greenhouse at 20–22 °C in soil under natural light conditions. Briefly, leaves were homogenized in ice-cold grinding medium A, containing 20 mM Tricine (pH 7.6), 0.4 M sorbitol (or 0.3 M NaCl [30]), 5 mM MgCl_2_ and 5 mM KCl, and filtered through six layers of medical gauze pads. After discarding the remaining debris by centrifugation at 200×*g* for 2 min, the supernatant was centrifuged for 5 min at 4000×*g* and the pellet was resuspended in 10 ml osmotic shock medium containing 20 mM Tricine (pH 7.6), 5 mM MgCl_2_ and 5 mM KCl. After a short, 5–10 s, osmotic shock, breaking the envelope membrane and allowing the replacement of the stroma liquid with the reaction medium, the osmolarity was returned to isotonic conditions by adding equal volume of double strength medium. This suspension was then centrifuged for 5 min at 4000×*g*. The thylakoid samples were stored at 4 °C until further treatments and/or use in the experiments.

### pH treatments

The thylakoid samples were washed twice with reaction medium A, then resuspended for thermoluminescence (TL) measurements in the same medium adjusted to different pH values (pH 7.5, 6.5, 5.5 or 4.5) or in the same, D_2_O-based medium, to p^2^H 8.0, 7.0, 6.0 and 5.0, for small-angle neutron scattering (SANS) experiments. The chlorophyll concentration was adjusted to 1–2 mg/ml for SANS measurements and 1.3 mg/ml for TL measurements. The pH-treated thylakoid membranes were kept in dark at room temperature for 30 min before the measurements. After 30 min, half of the samples were used in the measurement, and for the recovery experiments the remaining samples were washed twice with reaction medium A and then resuspended in the same medium at pH 7.5 or p^2^H 8.0 for the TL and SANS measurements, respectively, and were kept in dark at room temperature for 30 min.

### SANS experiments

SANS measurements were performed on the SANS-II small-angle neutron scattering instrument at Paul Scherrer Institute, Villigen, Switzerland, as previously described [29]. The wavelength, sample-to-detector distance and collimation were 6 Å, 6 m and 6 m, respectively. The isolated thylakoid membranes were measured at room temperature in a quartz cuvette of 2 mm optical path length in the presence of ∼0.4 T horizontal magnetic field with the field vector perpendicular to the neutron beam. The samples were measured for 2^*^5 min (with sorbitol as osmotic medium) and 5^*^2 min (with NaCl as osmotic medium).

### SANS data treatment and fitting procedures

All experimental data are normalized to the number of beam monitor counts; instrumental backgrounds and scattering from the suspending media were subtracted from the scattering profiles. The detector efficiency was calculated from background-subtracted water measurement. The primary data were treated with the Graphical Reduction and Analysis SANS Program for Matlab - GRASP (developed by Charles Dewhurst, ILL). The obtained two-dimensional data were reduced from 2D to 1D profile via radial or azimuthal averaging. The radial averaging was performed in two 75° sectors around each opposite Bragg diffraction peaks [31] in order to obtain intensity (I) versus scattering vector or momentumtransfer (q) curves.

The scattering curves were fitted with the phenomenological model expressed by the linear combination of a constant, power and Gauss functions in the q region of 0.01-0.033 Å^-1^ (sorbitol) and 0.015-0.042 Å^-1^ (NaCl) around the Bragg peak in order to determine the center position of the Bragg peak (q^*^) [31]. The center position of the Bragg peak was used to calculate the thylakoid membrane repeat distance (RD) values, according to RD=2π/q^*^. In order to better visualize the shift in the position of the Bragg peak we also used the Kratky-plot (*I*(q).q^2^ vs q) [32], where *I(q)* was obtained as follows: the radially averaged intensity in vertical orientation (with an opening angle of 75°) was subtracted from the radially averaged intensity in horizontal orientation (with an opening angle of 75°).

In order to provide information on variations in the mosaicity of membranes, we determined *I*(φ), the angular dependency of the scattering intensity. To this end, 2D SANS profiles were azimuthally integrated across the q region of 0.017-0.44 (sorbitol) and 0.025-0.040 (NaCl) Å^-1^ for 360° interval with 5 pixel binning (φ is the azimuthal angle).

The I(φ) curves were fitted with the sum of a constant and a Lorentzian function; its widths (FWHM, full width at half maximum) provides information on extent of the magnetic orientability of sample, and thus on its anisotropy.

A quantitative characterization of the degree of orientation can be obtained by Hermans orientation function [33], which takes value of 1 or -0.5 when the membranes are completely oriented parallel or perpendicular to the direction of reference, respectively, and 0 for the case of random orientation. In our case, the direction of reference is the direction of magnetic field, and for perfectly aligned sample the value would be 1.0.

### Thermoluminescence measurements

The measurements were carried out using a home-built thermoluminescence apparatus [34]. A single-turnover saturating flash excitation was applied at -30 °C; the heating rate was 20 °C/min [35]. These measurements were used to control the efficiency of our low pH treatments. The observed low-pH induced reversible shifts of the B-band (data not shown) were in perfect agreement with literature data [36, 37].

## Results and discussion

As reported earlier [38] the SANS signal of magnetically oriented thylakoid membranes is dominated by well discernible scattering peaks with maxima on the 2D image in the direction parallel with the direction of the applied magnetic field (Figure 1A). Upon acidification of the suspension medium the observed diffraction peak became more flat and the Bragg peaks largely diminished (Figure 1B). Resuspending the low-pH treated thylakoid membranes in the p^2^H 8.0 medium largely restored the original 2D profile (Figure 1C). It is interesting to note that these low-pH induced variations in the 2D SANS profiles closely resemble the ΔpH-dependent light-induced changes in the 2D scattering profiles of isolated thylakoid membranes [29, 31, 38].

**Figure 1:**
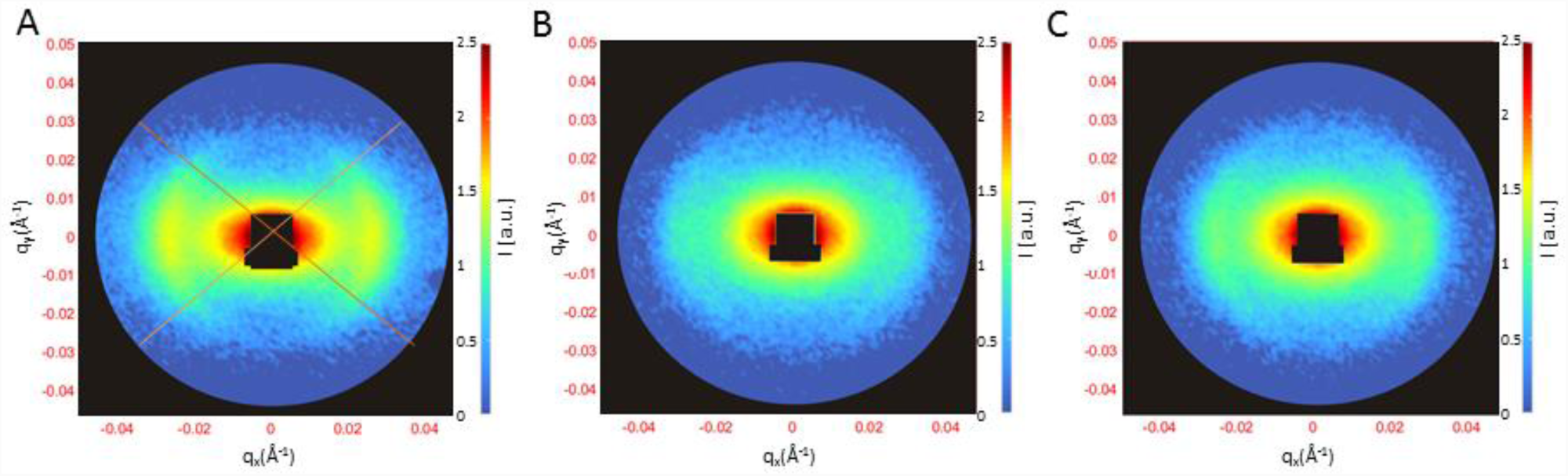
2D small-angle neutron scattering profiles of magnetically oriented thylakoid membranes isolated from pea leaves and suspended in a p^2^H 8.0 reaction medium (A), resuspended in the same medium adjusted to p^2^H 5.0 (B), and returned to the p^2^H 8.0 medium (C).

In order to obtain quantitative information about the low-pH induced reorganization of the thylakoid membranes we performed sectorial averaging of the 2D scattering curves – allowing precise determination of the diffraction peak (and hence the RD of the thylakoid membranes), and also investigated the angular dependency of the 2D scattering signal around the diffraction peak. In earlier studies, we have shown that the osmoticum used in the reaction medium significantly influences the structure of the isolated thylakoid membranes, and that NaCl retains much better the in-vivo structure of the thylakoid membranes than sorbitol [29]. For this reason, we performed the experiments both in sorbitol- and NaCl-based media.

The radially averaged scattering curves revealed similar and strong influence of the acidity of the suspension medium on the multilamellar arrangement of the thylakoid membranes for the two types of reaction media (Figure 2A and B). In both cases, the diffraction peak around 0.019 Å^-1^ (sorbitol) and 0.027 Å^-1^ (NaCl) was shifted towards higher momentumtransfer values while its intensity was diminished. These variations are best seen using Kratky plots of the data (insets in Figure 2).

**Figure 2.**
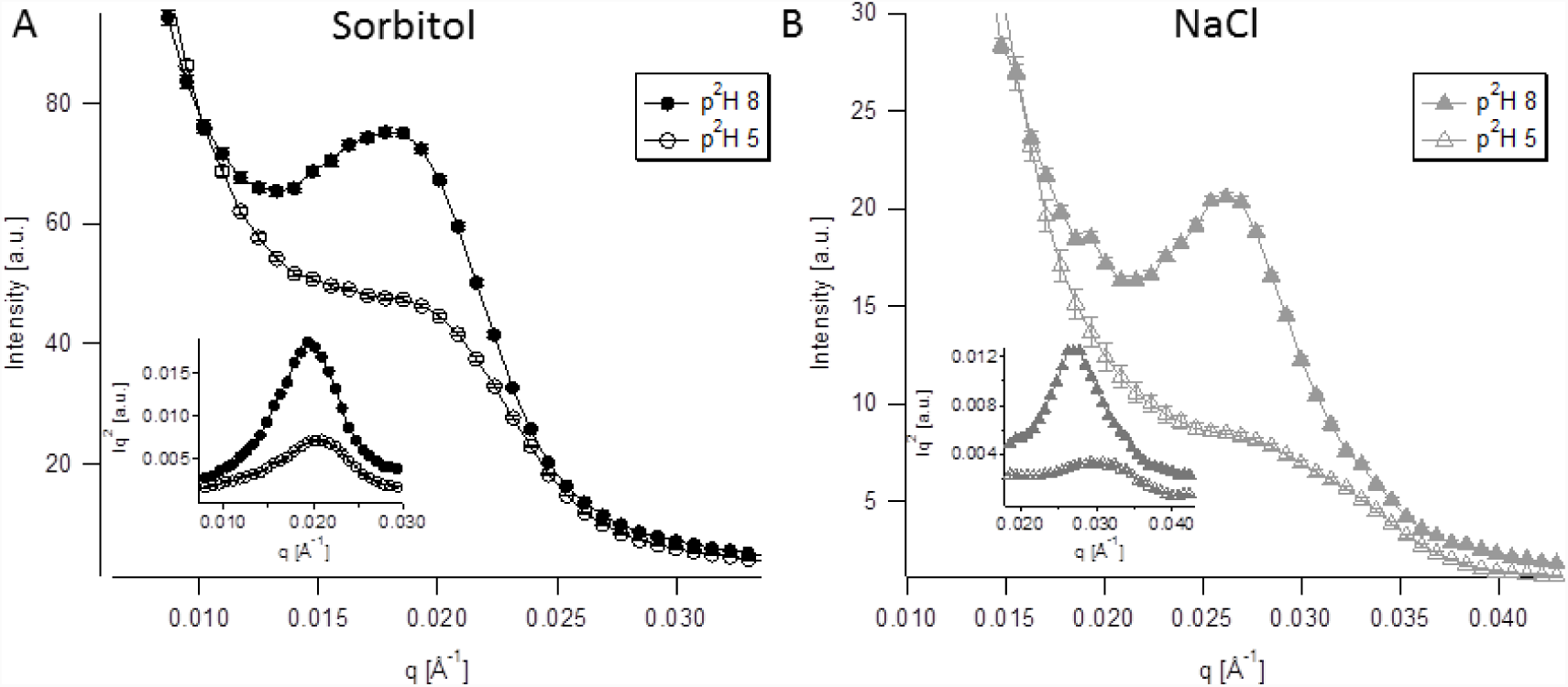
Sectorially averaged scattering curves of the thylakoid membranes suspended in p^2^H 8.0 and p^2^H 5.0. The suspension medium contained, as osmoticum, sorbitol (A) or NaCl (B). Insets are the Kratky plot of the same data [32].

We determined the center position of the diffraction peaks and calculated the average RD of the thylakoid membranes at various p^2^H conditions (Figure 3A). Upon acidification the RD was observed to decrease, while upon resuspension in the original medium (p^2^H 8.0), the original RD values were almost perfectly recovered. This acidification-induced reversible shrinkage of the thylakoid membranes strongly resembles the effect of illumination, observed earlier on isolated thylakoid membranes [29, 31, 38, 39]. Similarly to the case of light-induced variations observed on the radially averaged SANS curves of thylakoid membranes the intensity of the fitted Gaussians was reduced also upon acidification (see Figure 3B) – indicating a disorder in the periodic membrane ultrastructure; these changes were, however, only partly reversible upon resuspension in media with p^2^H 8.0, especially after exposures to p^2^H 5.0.

**Figure 3.**
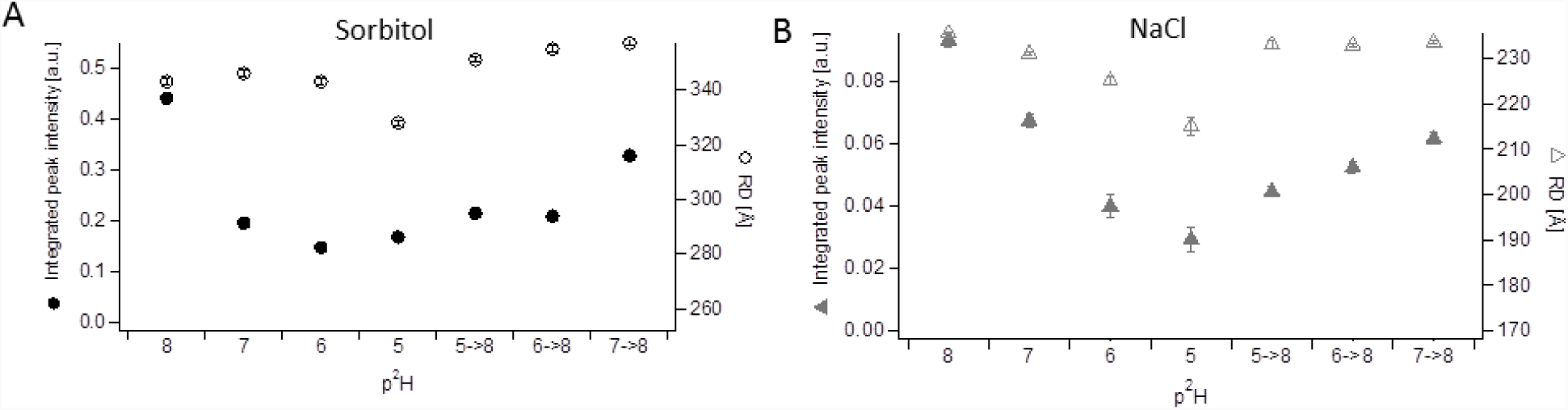
Physical parameters of Bragg peaks at various p^2^H conditions: the calculated average RD and the integrated Bragg peak intensity (B) in sorbitol-containing medium (A) and in NaCl-containing medium (B). The error bars show the standard errors of the RD and the integrated peak intensity values and originate from the uncertainty of the fitting.

As concerns the low-pH induced disorder, the analysis of the angular dependence of the SANS signal also reveals significant changes. As shown in Figure 4, upon acidification the orientability of the multilamellar membranes was significantly reduced. For the interpretation of these changes we discuss below the origin of the broadening of the Bragg peak, including the case of thylakoid membranes suspended in p^2^H 8.0 media.

**Figure 4.**
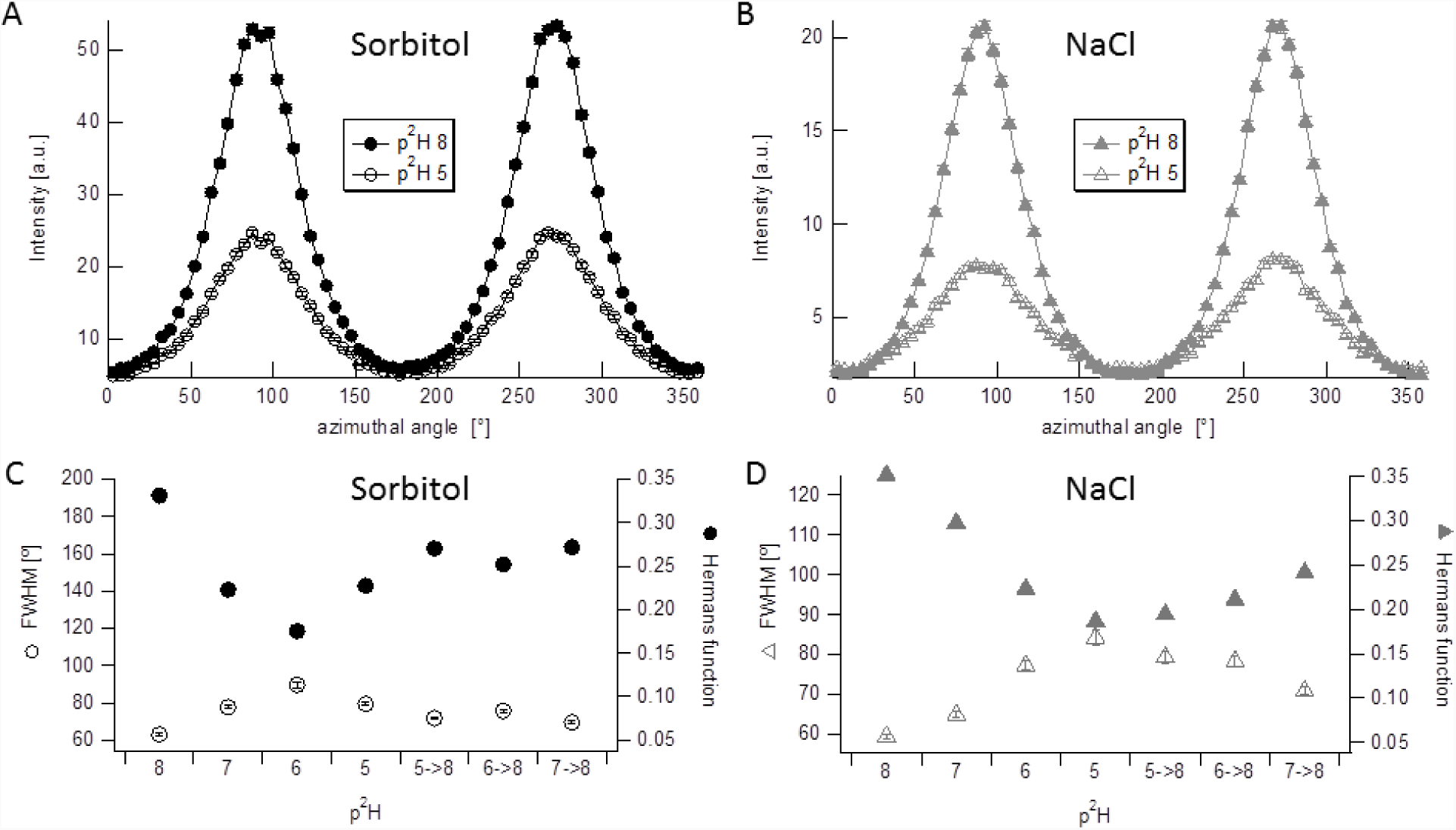
The angular dependences of the scattering intensity (A,B) of the thylakoid membranes suspended in p^2^H 8.0 and p^2^H 5.0 with the suspension media containing, as osmoticum, sorbitol (A) or NaCl (B); and the dependences of the full width at half maxima (FWHMs) of the Lorentz function and the Hermans’ orientation function on the p^2^H treatments of the thylakoid membranes with the suspension media containing, as osmoticum, sorbitol (C) or NaCl (D). The error bars in B and C show the standard errors of FWHM.

In an external magnetic field intact thylakoid membranes tend to align perpendicular to the magnetic field. However, there are irregularities in the membrane system, and the chloroplasts and membranes often assume banana-shape. Therefore, not all grana sections, which give rise to the scattering peak, can be oriented exactly perpendicular to the magnetic field and the majority of membrane normals will have a non-zero angle relative to the magnetic field. This leads to a relatively broad distribution of grana in Bragg condition [31]. The scattering signal from these imperfectly oriented grana will exhibit, for symmetry reasons, two Bragg peaks with a broad spread in azimuthal angles around the horizontal direction. Additional factors, which contribute to the broadening of the scattering peak, the finite spectral bandwidth of the monochromatized neutron beam and other experimental factors, such as the detector resolution and the finite size of the incident beam, that contribute to the smearing of the Bragg peak can be neglected since (i) their contributions are small and (ii) they do not differ sizeably for samples before and after the low-p^2^H treatments.

Variations in the angular dependence of the scattering signal, the reduced orientability of the multilamellar membranes upon acidification suggest an increased mosaicity, i.e., an increase in the spread of membrane plane orientations.

Based on the above data it can be concluded that the observed low-p^2^H induced smearing and broadening of the Bragg peak and the increased mosaicity of the membranes reflects a loosening in the periodic order of the thylakoid membranes that evidently arises from some undulations or membrane bending. These membrane reorganizations, along with the low-p^2^H induced shrinkage, might be related to the lateral rearrangements of the protein complexes that are thought to be responsible for the observed changes in the chiral macrodomains (i.e., in the psi-type CD) and in the distribution of absorbed excitation energy between the two photosystems – regulated by PsbS [24].

In general, these results, in line with similar observations [29, 38, 40-42] underline the remarkable flexibility of the thylakoid membrane ultrastructure, which should thus not be portrayed as simply providing a scaffold for the photosynthetic functions but also actively participating in the energy conversion steps and in different regulatory functions.

## Acknowledgement

This work is based on experiments performed on the SANS-II beam-line at the Swiss spallation neutron source SINQ, Paul Scherrer Institut, Villigen, Switzerland. This work was supported by grants from the Hungarian Scientific Research Fund (NKFIH OTKA K 112688, GINOP-2.3.2-15-2016-00001 to GG) and Scientific Exchange Program-NMS.CH (SCIEX) (Project number 13.098 to RÜ).

